# Raman microspectroscopy imaging analysis of extracellular vesicles (EVs) biogenesis by filamentous fungus *Penicilium chrysogenum*

**DOI:** 10.1101/2021.11.04.467387

**Authors:** Ashok Zachariah Samuel, Shumpei Horii, Takuji Nakashima, Naoko Shibata, Masahiro Ando, Haruko Takeyama

**Author notes:** **Corresponding author** Correspondence to Haruko Takeyama.

## Abstract

Mechanism of production of extracellular vesicles (EVs) and their molecular contents are of great interest owing to their diverse roles in biological systems and are far from being completely understood. Even though, cellular cargo release mediated by EVs have been demonstrated in several cases, their role in secondary metabolite production and release remains elusive. In this study we investigate this aspect in detail using Raman micro-spectroscopic imaging. We provide considerable evidence to suggest that the release of antibiotic penicillin by filamentous fungus *Penicillium chrysogenum* involves EVs. Morphological modifications of the fungal body during biogenesis, changes in cell composition at the locus of biogenesis, and major molecular contents of the released EVs are also revealed in this study.

**Importance:** Extracellular vesicles (EVs) play a key role in cellular communications. EVs role in functioning of fungi are relatively less explored. Here we show selective enrichment of chemical contents at certain locations of mycelium of *P. chrysogenum* forming protruding regions. Secondary metabolite penicillin is excessively localized in them. We provide evidence to show that EVs are released from these protrusions. Raman imaging has been applied for molecular profiling of the mycelium and for characterizing chemical contents of the EVs. Our study suggests a possible general role of EVs in the release of antibiotics from the producing organisms.

## Introduction

All cellular systems including microorganisms, multicellular organisms, and even circulating cells in the body fluids of higher organisms, require frequent transmembrane molecular transport to function effectively. The secretion of the molecules in vesicular cargo has received renewed interest originating from their roles in cellular communications (1), viral/bacterial infections (2, 3), immune system response (4, 5), cancer (6, 7) etc. Initial studies have attributed the origin of such vesicular *cargo* units to be mainly endosomal in nature (8). Recently, plasma membrane mediated generation of extracellular vesicles (EVs), with sizes varying from 50 nm to 1,000 nm, have been observed in several living systems (2, 9). EVs’ role in pathogenesis is becoming more evident lately, for instance, EVs produced by infectious *Staphylococcus aureus* delivers the toxin to human host cells (10). Evidence support that bacterial probiotic effects are also mediated through EVs containing effector molecules (e.g. *Lactobacillus casei*) (11). Inspired by EVs roles in living systems, EVs based therapeutic strategies are also being developed for RNA and protein delivery, and targeted drug delivery (12). Addressing the mechanism of production and the molecular constitution of EVs produced by microorganisms are especially important because of their complex roles in pathogenesis (13) to inventive roles in therapeutics (14).

Mechanism of production of EVs and their functional roles in diverse cellular systems are currently being actively investigated. According to the current understanding, in gram negative bacteria, budding (15) of the outer membrane occurs at the regions having weaker crosslinking with the peptidoglycan (PG) layer (16), and then the EVs detach from the cells without compromising the membrane integrity (17). The degradation of PG layer by a biological agent can also result in explosive cell lysis leading to generation of a large number of EVs (18). In bacteria *Pseudomonas aeruginosa* and *Bacillus subtilis* the role of endolysins in effecting PG degradation leading to the production of EVs has been demonstrated (18–21). In the case of filamentous fungi, with thicker polysaccharide cell wall and without phospholipid outer membrane, the mechanism of EVs biogenesis remains unclear. Owing to the presence of thick polysaccharide cell wall, with inherent porosity in the range of a few nanometers (19), a larger opening in the cell wall might be necessary for EVs secretion. *Cryptococcus neoformans* produces virulent polysaccharide glucuronoxylomannan in Golgi secreted vesicles and transport it across cell wall (22). Such fungal EVs secretion has also been demonstrated in *Histoplasma capsulatum*, *Paracoccidioides brasiliensis*, and *Aspergillus fumigatus* (23), however, no cell wall damage/opening has been observed in these cases. Despite diverse mechanisms involved, there is considerable evidence to suggest that EVs generation is a well-regulated biological process (24– 27) rather than a stochastic event occurring at leaky regions of the membrane.

Since EVs are biologically regulated depending on the functional requirements of the organism, existence of subpopulations of EVs with varying sizes (28), composition (29–31), and contents (32), are possible. However, such properties are rather difficult to explore (24–26). Secondary metabolites produced by microorganisms, including several fungi, are central in human defense against pathogenic infections (33). The production, localization, and the release of these key molecular secretions from the cell body are of considerable interest particularly because of antibiotic resistance prevalent in microbial community (34, 35). It is unclear if the release of secondary metabolites involves EVs. In this study, we investigate the possible role of EVs in the release of penicillin produced by *P. chrysogenum*. For a detailed study, we have employed molecular structure specific Raman micro-spectroscopic imaging technique in conjunction with multivariate curve resolution (MCR) analysis. Since the antibiotic molecule, penicillin G, is structurally different from common cellular constituents, its localization and a possible release involving EVs can be investigated with this technique. The effectiveness of this methodology has been demonstrated in our recent study by performing molecular profiling of lipid droplets in human liver cells (36). Our modified MCR analysis routine (36, 37) quantitatively decomposes highly overlapped composite Raman spectral data into spectra of biomolecules revealing their individual spatial distributions. Owing to the presence of thick polysaccharide layer at the cell boundaries, strong Raman signal specific to polysaccharides can be obtained (38), which will then allow inspecting the integrity of the cell wall in case EVs mediate the release of the antibiotic. The exquisite ability of Raman imaging technique to detect production and localization of penicillin inside mycelia of *P. chrysogenum* has been demonstrated in our recent study (38). Here we investigate the penicillin release by EVs biogenesis from *P. chrysogenum*.

## Results

Penicillin production in the *P. chrysogenum* culture extract was confirmed with antimicrobial activity test (inhibition zone 50 mm/10 μl per disk) and LC-MS analysis (Fig. S1). Raman imaging was conducted on the filamentous mycelium and MCR data analysis (*see* materials and methods) was performed. Results of the MCR analysis are given in Fig. 1. Characteristic spectra corresponding to proteins, lipids, cytochrome c, penicillin, polyphosphate and polysaccharide can be accurately identified (For details of spectral assignments *see* materials and methods: spectral assignments). Raman images showing spatial distribution of these biomolecules in the mycelium are provided in Fig. 1b. In the Raman images, proteins and lipids can be seen more or less continuously distributed throughout the cell. Polyphosphate and cytochrome c are distributed at certain regions of the cell, in agreement with our previous study (38). A protrusion originating from mycelium is indicated with a circle. An overlay of proteins and polysaccharides (Fig. 1c), and a drawing indicating protrusion (Fig. 1d) are also shown. A high concentration of penicillin can be noticed at the protrusion (Fig. 1b).

**FIG. 1.**
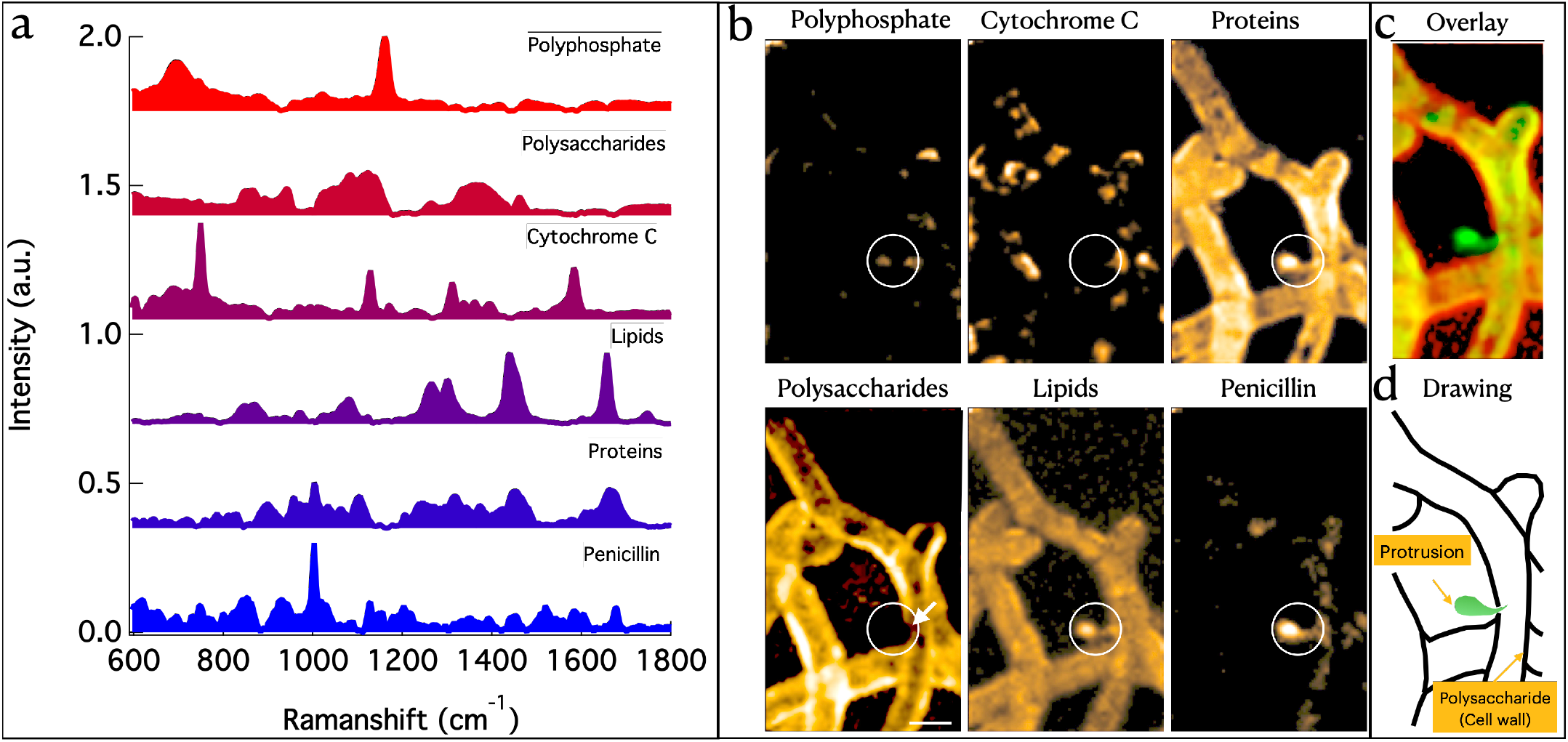
EVs budding. a) MCR separated Raman spectral components. A full list is provided in the Fig. S2. b) Raman images of chemical constituents of *P. chrysogenum* filamentous body. The protrusion from the cell body is circled. An apparent location of a small opening in the cell wall is indicated with a white arrow. Size of the small opening cannot be accurately estimated (apparently <<1μm) from the image due to the limited spatial resolution of the technique. Scale bar 5 μm. c) Overlay of polysaccharide and proteins images: red color – polysaccharide and green color – proteins. d) A drawing explaining the observations. The black line indicates polysaccharide cell wall, and the protrusion is shown in green color.

We examined the culture medium to confirm the presence of EVs. Several large (∼1 μm) EVs were found in the culture medium outside the cell regions (Fig. 2a). It is important to examine the composition of the EVs in order to elucidate the possible role of penicillin in their biogenesis. Raman images of the EVs (Fig. 2a; Fig. S3) and the results of MCR analysis (Fig. 2b) are provided in Fig. 2 a-c. Proteins, penicillin, and carotenoid were detected inside EVs (Fig. 2b), and on an average these EVs contain ∼4 ± 1 mM protein, ∼38 ± 19 mM penicillin and ∼5 ± 5 μM carotenoid (Fig. 2c; all concentrations estimated relative to Raman intensities of standard compounds in solution. For details see materials and methods; ± std. dev.; carotenoid is resonance enhanced). No size dependent variation in penicillin content was observed in the EVs (Fig. 2d). It is important to note that six different molecules were detected in the filamentous cell body, however, only protein, penicillin and carotenoid were detected inside the EVs. Carotenoids were detected in the EVs and not in the main filamentous body of the organism (38). Resonance enhancement makes it possible to detect carotenoid even at very low concentrations (μM) using Raman spectroscopy. Based on the observed peak positions, the carotenoid spectral component has been assigned to β-carotene (39). However, several different carotenoids give Raman spectra that are only marginally different from one another and exhibit peak shift associated with mode of binding in biological systems (39). HPLC analysis of the concentrated extract from the *P. chrysogenum* culture (supernatant containing EVs) was performed and the presence of β-carotene was confirmed (Fig. 2e). Notably, polysaccharide is absent in the EVs (Fig. 1) suggesting these are not broken fragments of the main filamentous body. Owing to the relatively lower concentration of lipids in the EVs membrane, it was not detected in the Raman analysis. In order to verify the presence of lipid membrane surrounding the EVs, we performed staining with a membrane selective dye (FM 4-64; (N-(3-Triethylammoniumpropyl)-4-(6-(4-(Diethylamino) Phenyl) Hexatrienyl) Pyridinium Dibromide)). As expected, the dye has labelled the EVs, which affirms its lipid membrane bound structure (Fig. 2f).

**FIG. 2.**
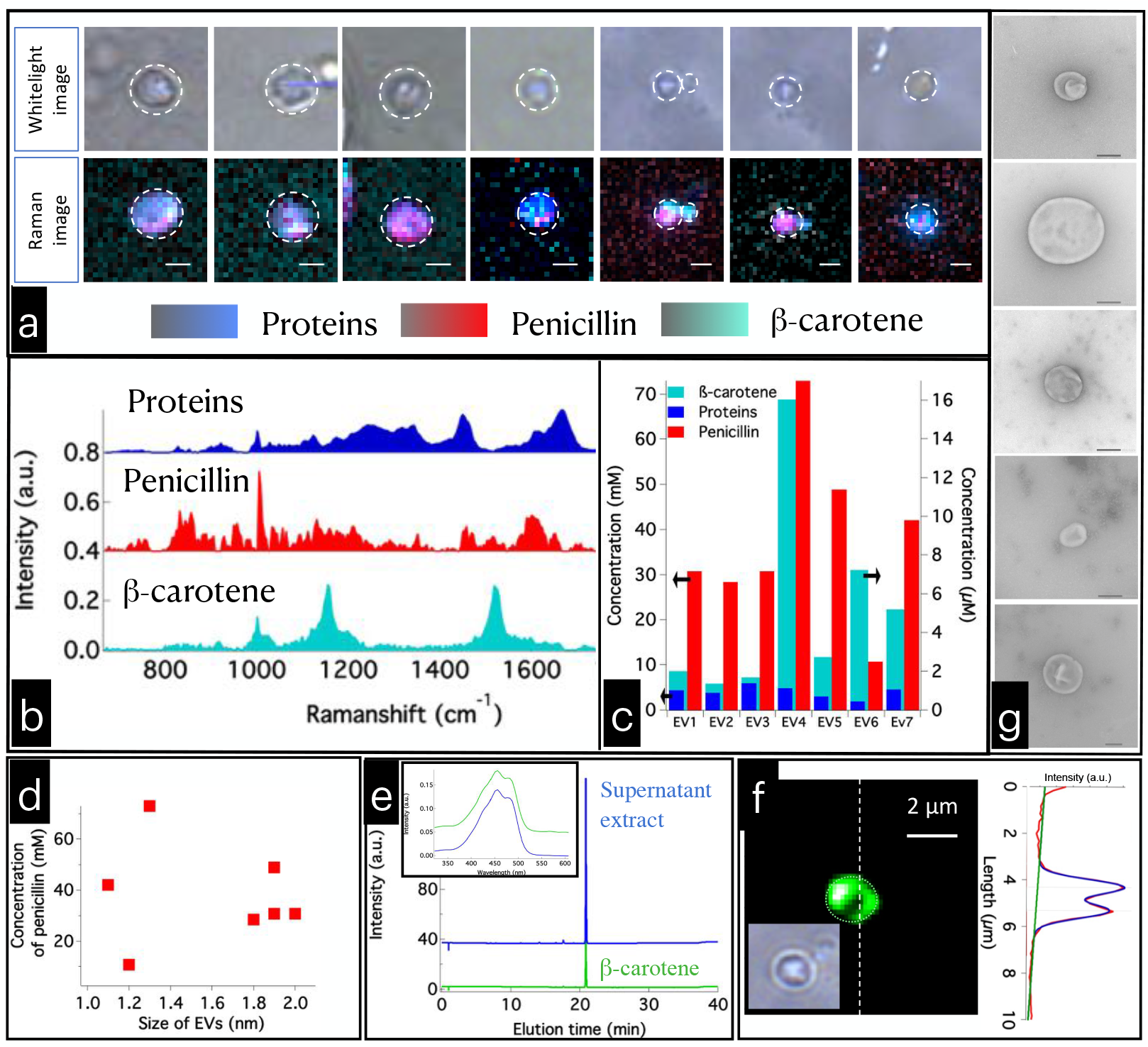
EVs observation and their characterization. a) white light and Raman images (overlay of β-carotene, proteins, and penicillin Raman images; for individual images *see* Fig. S3) of EVs from *P. chrysogenum* observed in the culture. Scale bar 1 μm. b) molecular constituents of the EVs imaged with Raman microspectroscopy. Molecular compositions of the EVs are characteristically different from that of filamentous body of the organism. c) estimated concentrations of molecular contents of the EVs. d) Variation of penicillin content depending on the size of EVs. e) Results of HPLC analysis of concentrated supernatant containing EVs separated from 10 duplicate *P. chrysogenum* cultures. Elution profiles obtained by monitoring at 450 nm wavelength. UV absorption spectra of standard β-carotene solution, and β-carotene in supernatant extract (inset). f) Fluorescence image of an EVs stained with FM 4-64 membrane selective dye. Intensity profile along the line indicated in the fluorescence image is also shown. Fluorescence imaging was performed with the same Raman microscope used in the study. g) TEM images of the EVs. Scale bar 200 nm.

TEM images of the EVs from the supernatant of the culture broth are shown in Fig. 2g. Lighter EVs can be separated from denser cell debris by performing density gradient ultracentrifugation (40). LC-MS analysis was performed to examine the presence of penicillin in different layers of the sample after ultracentrifugation. Significant amount of penicillin was detected in the top two layers (each 2 ml), while penicillin was not detected in the subsequent layers (Fig. S4). We have performed TEM analysis of these layers and a large number of EVs were detected in the first two layers (Size: 180 nm to 600 nm) while cell debris were seen in the subsequent layers (Fig. S4). This result reaffirms the production of EVs containing penicillin by *P. chrysogenum*.

Our attempts to obtain Raman images of EVs after ultracentrifugation was not successful due to fortuitous ingress of iodixanol (41) into the isolated EVs (Fig. S5). This method must be modified in future work.

*P. chrysogenum* is known to produce spores of sizes comparable to that of EVs observed and therefore it is necessary to distinguish the EVs from spores. We have separately cultured spores of *P. chrysogenum* and Raman imaging was conducted. The results of MCR analysis indicated prominent polysaccharide (42) signature (Fig. S6), which was absent in the vesicular structures.

Ca^2+^ is a chemical constituent keeping polysaccharide layer intact in the fungus cell wall. Removal of Ca^2+^ by adding small amounts of EDTA should weaken the cell wall and thereby it may facilitate the formation of protruding regions (43). We have therefore added EDTA into the culture medium to facilitate the EVs production. As expected, several protrusions were observed in the filamentous body (Fig. S7). The results of the Raman-MCR analysis are given in Fig. 3.

**FIG. 3.**
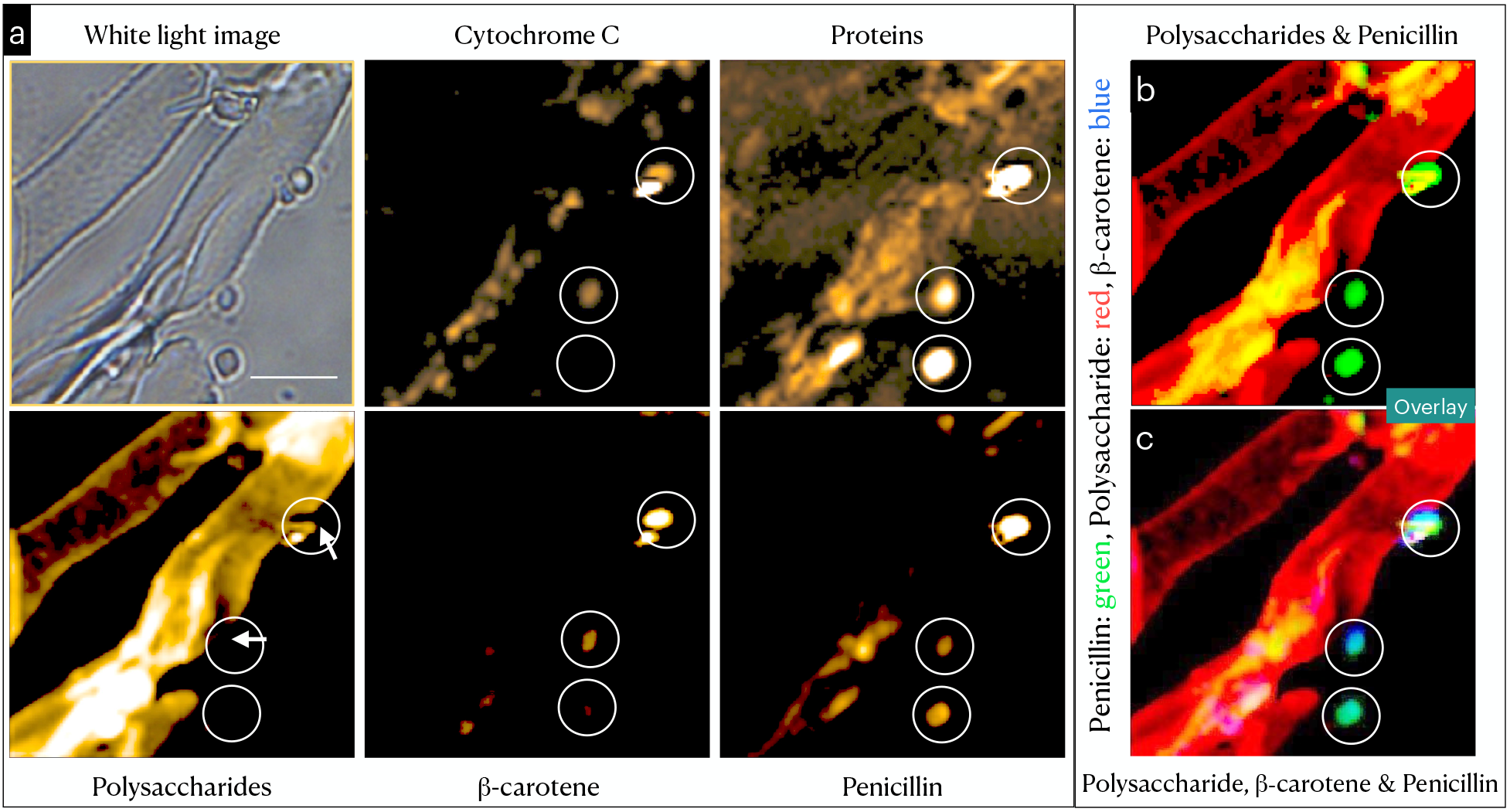
Biogenesis of EVs. a) Raman images of EDTA treated *P. chrysogenum*. Protrusions from the cell wall can be seen at several regions, and all of them have high concentration of penicillin. b) An overlay of polysaccharides and penicillin; it indicates the absence of polysaccharides in the protruding regions. c) An overlay of polysaccharide, β-carotene and penicillin: Apparently, β-carotene is present exclusively in the protrusions. Scale bar 5 μm. For the corresponding spectral components, overlay with white light image, Raman image of lipids etc. *see* fig. S7.

Appearance of several protruding regions in the filamentous body suggests that addition of EDTA has facilitated its formation and therefore the biogenesis of the EVs. All these protrusions were rich in penicillin, similar to the protrusions observed in the culture without EDTA. Raman images given in Fig. 3 indicate *β*-carotene as one of the constituents of the protrusions (except one). Notably, *β*-carotene was not seen as a chemical constituent of the filamentous body. Additionally, *β*-carotene was a constituent of all the EVs shown in Fig. 2a.

## Discussion

The appearance of protrusions in the main filamentous body with characteristically different chemical composition is clearly demonstrated with Raman imaging (Fig. 1b). A comparison of the penicillin and polysaccharide distributions reveals interesting aspects. Penicillin is highly concentrated in the balloon like protrusion (Fig. 1b: penicillin) compared to the main filamentous body (in terms of Raman intensities, ∼18% in the protrusion compared to ∼4% in the mycelium (Fig. 1b)). Interestingly, the protrusion does not have polysaccharide (Fig. 1b: polysaccharide) boundary, and where it originates, the polysaccharide cell wall appears comparatively thin and, apparently, with a small opening (Fig. S7e). Endolysin induced hole formation in the PG layer and consequent EVs release is known to occur (19). However, in the present case no external chemical inducer was employed. If the opening was a physically formed, then there is no reason for penicillin concentration at the protrusion to be considerably different from cytoplasmic composition. A characteristically high concentration of penicillin, we believe, implies a molecular mechanism that involves penicillin itself, and probably an urge to release it. The following scenario is possible. In order to release penicillin, it is first accumulated at specific regions of the filamentous body. A protrusion is then formed through the cell wall, from which, probably, EVs containing penicillin are subsequently released. Supporting this hypothesis, we detected several EVs containing penicillin in the culture supernatant (Fig. 2a and g) with high relative composition of penicillin is observed in the EVs similar to that in the protrusion. Moreover, no polysaccharides were detected in the EVs (hence not spores or fragments of the mycelium) and a lipid selective fluorescent dye has labelled the EVs. Relatively high concentration of penicillin in both the protrusions and the isolated EVs indicates that the production of EVs is linked to the formation of the protrusions.

EDTA addition has facilitated the formation of protrusions suggesting that a higher porosity (or an opening) in the cell wall is necessary for the biogenesis (EDTA weakens cell wall(43)). Further, β-carotene is observed in the EVs and in the protrusions but not in the other parts of mycelium (Fig. 3). Therefore, most probably *β*-carotene is produced during the biogenesis of the EVs to protect them from possible environmental stress (44, 45). Penicillium sp. is known to produce carotenoid under oxidative stress (46). Production of *β*-carotene at the protruding regions also supports the fact that the observed event is, most probably, the biogenesis of the EVs. It is also interesting to note that all the protrusions do not have exactly same composition (compare intensities of the circled regions in Fig. 3). In Fig. 3, cytochrome c is seen in two of the protruding regions in spite of their absence in isolated EVs. β-carotene was also absent in one of the three protrusions (Fig. 3). In the absence of additional evidence, we attribute it to the different stages of biogenesis. Absence of β-carotene in the protrusion in Fig. 1 could also be reasoned along the similar lines. Further, we find no evidence to suggest that EDTA induces excess β-carotene production in *P. chrysogenum*.

High penicillin content in the EVs is evident from the Raman images (Fig. 2), which is also supported by LC-MS results (penicillin was detected only in the layers containing EVs; Fig. S6). Taken together, it is clear that the EVs containing relatively larger amount of penicillin are produced by *P. chrysogenum*. During different stages of biogenesis, the protruding regions can have different compositions. The contents of the protrusions are, apparently, progressively regulated, and EVs with similar molecular contents are eventually released by the organism.

Penicillin biosynthesis by *P. chrysogenum* involves multiple enzymatic reactions: a) synthesis of ∂-(L-α-amino adipyl)-L-cysteinyl-D-valine (ACV) tripeptide mediated by ACV synthetase, b) ACV is converted to isopenicillin (IPN) by Isopenicillin N synthase (IPNS) and finally, c) enzymatic conversion of isopenicillin intermediate to penicillin G, either completed in a single enzymatic step involving acyl CoA:IPN acyltransferase, or through a two-step enzymatic reaction involving an intermediate 6-aminopenicillanic acid (6-APA) (47, 48). The final step (i.e., step c) of penicillin biosynthesis occurs in specific locations called microbodies (peroxisomes) (48–50), which are of dimensions comparable to the EVs observed in our study (49, 51). Interestingly, increased number of peroxisomes inside filamentous body of *P. chrysogenum* has also been correlated with higher penicillin production (50, 52). It is highly probable that the microbodies are one of the major sources of the vesicular (EVs) contents produced by *P. chrysogenum*.

Penicillin release by exocytosis mediated by vacuoles produced by pexophagy of microbodies has been proposed in an earlier study based on electron microscopy images (53). In spite of the difference in the suggested mechanism, our conjecture of relation between contents of microbody and EVs is supported by this study. EVs with heterogeneous molecular contents secreted by fungus *Cryptococcus neoformans* were earlier observed in the supernatants of the cultures (54). It was hypothesized that these EVs are Golgi synthesized and later brought to near plasma membrane. The mechanism of cell wall passage, however, was not clear. Our results show the generation of EVs though a process involving extrusion of specific cytoplasmic contents though the cell wall, albeit in a different fungus.

Raman spectroscopy, being a molecule specific analytical method, combined with our novel MCR routine (36, 37) allowed detection of multiple chemical species inside the filamentous body of *P. chrysogenum*. This exquisite ability has been utilized in the present study to address several important questions concerning the mechanism of biogenesis, and the chemical composition of the EVs. Our study provides first consolidated evidence for the release of secondary metabolite - penicillin in the present case - from a fungus mediated by EVs. We show that penicillin is enriched at the regions of the fungal body where the EVs biogenesis occur. Through the polysaccharide cell wall penicillin rich protrusions form, and the contents are then released in the form of EVs. However, it remains unclear from our study whether this is the only mechanism for the release of penicillin. Our findings and suggested mechanism are also supported by the recent study by Toyofuku et al., albeit in Gram-positive bacteria (19). We believe that penicillin accumulation, polysaccharide wall ‘opening’ and release of EVs are all steps in the major cascade in the biogenesis of the EVs in *P. chrysogenum*. It is possible that EVs also, in general, mediate the release of secondary metabolites from microorganisms.

## Materials and Methods

### Raman Spectroscopy

Raman microspectroscopic imaging measurements were carried out with a laboratory-built confocal Raman microspectrometer. A 532 nm output of a Nd:YAG laser (Compass 315M; Coherent Inc., Santa Clara, CA, USA) was used as the laser source. The laser beam was focused with a 100X (1.4 NA) objective lens (Plan Apo VC; Nikon Corporation, Tokyo, Japan) mounted on an inverted microscope (ECLIPSE Ti; Nikon Corporation, Tokyo, Japan). The back-scattered Raman light was measured with a spectrometer (MS3504i, 600 lines/mm; SOL Instruments, Ltd., Minsk, Republic of Belarus) and a CCD detector (Newton DU920-M; Andor Technology Plc., Antrim, UK). The laser power at the sample was ∼10 mW. A piezoelectric stage (custom-made; Physik Instrumente GmbH & Co. KG, Karlsruhe, Germany) was used to carry out Raman imaging (0.3 μm step size) with 1 sec exposure for each Raman spectral acquisition. Spectra were calibrated using indene. Raman data analysis and detailed image processing were performed with codes written in *Igor Pro*. Image J was also used to plot Raman images.

### MCR-ALS and SVD

Multivariate curve resolution (MCR) by alternating least squares (ALS) is performed by solving eqn. 1. Raman data matrix (A_m,n_) can be decomposed into spectral components (W_m,k_) and their concentration profiles (H_k,n_) (36, 37, 55–59).

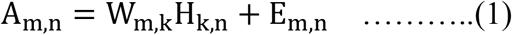

E is the residual. In the present case, A_m,n_ consists of n spectra from different spatial locations of the mycelium each with m wavenumber points. MCR-ALS is performed iteratively by minimizing the Frobenius norm ||A - WH||^2^ until negligible residual E results. Non-negativity constraints W ≥ 0 and H ≥ 0 are applied during the minimization procedure to obtain interpretable solutions because all the Raman spectral intensities and the concentrations are necessarily positive. The value of ‘k’ was guessed at first and later optimized. The number of independent spectral components with dominant singular values in singular value decomposition (SVD) served as an initial guess for ‘k’. Sparser MCR solutions were sought, when required, by introducing regularization schemes such as L1 norm (Lasso regression; α) or L2 norm (Ridge regression; β). L1 norm can be applied to H matrix or W matrix depending on situation as given below.

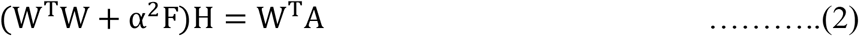

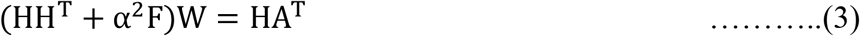

Where F is a k×k matrix in which all its elements are unity.

L2 norm can similarly be applied to H matrix or W matrix depending on situation as follows.

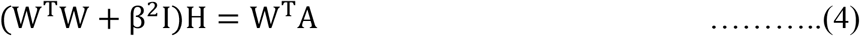

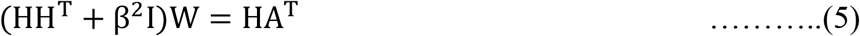

Where I is an identity matrix.

The optimized H matrix was plotted as a 2D Raman images.

### Cell culture

*P. chrysogenum* KF-425 was used as a fungus system producing antibiotics. Spores of strain KF-425 was inoculated into 10 mL preculture medium, containing 0.5% polypeptone, 2% glucose, 0.2% yeast extract, 0.1% KH_2_PO_4_, and 0.05% MgSO_4_·7H_2_O. It was incubated at 27 °C with shaking at 200 rpm for 2 days. About 1 ml of this pre-culture was then homogenized and inoculated into 100 mL main culture (F7; penicillin producing) medium, consisting of 2% sucrose, 1% glucose, 3% corn steep powder, 0.5% meat extract, 0.05% MgSO_4_·7H_2_O, 0.1% KH_2_PO_4_, and 0.3% CaCO_3_ (adjusted to pH ∼7.3 before sterilization). The liquid culture was incubated at 27 °C with shaking at 200 rpm for 5 to 6 days. After this, 1 mL of the culture broth was taken, 1 ml ethanol was added, and centrifuged at about 10000 rpm. HPLC and LC-MS experiments were conducted using the supernatant to confirm penicillin production. An aqueous solution of benzylpenicillin potassium (Penicillin G) (Fujifilm Wako Pure Chemical Co., Tokyo, Japan) was used as the reference compound. An antimicrobial activity test (inhibition zone assay) was also performed, as reported earlier (38), to confirm penicillin production.

EDTA (Ethylenediaminetetraacetic acid) experiment: On the third day of *P. chrysogenum* culture, 2 ml culture was separated into a new test tube and 100 μL of 0.5M EDTA solution was added, and culturing was continued for another 2 days. Afterwards filamentous *P. chrysogenum* cells were imaged with Raman microspectroscopy

### Antibacterial Activity Test

Antibacterial activity test was conducted using Kocuria rhizophira NBRC 103217. The antimicrobial activity was evaluated by a disk-diffusion agar method using paper disks (φ 8 mm) soaked with 10 μL of *P. chrysogenum* culture extract. The paper disc was placed on an agar plate seeded with *K. rhizophira*. The agar plate was then incubated.

### Transmission electron microscopy (TEM)

TEM images were recorded using JEM-1400 Flash Electron Microscope. For TEM imaging, the P. chrysogenum culture was centrifuged at 3500 rpm, filtered through 1 μm membrane filter to remove cells and other large debris. The samples were then negatively stained with 1% uranyl acetate solution and dried on the copper grid. Different layers after ultracentrifugation were also used as samples for detecting EVs.

### Gradient ultracentrifugation

Iodixanol solution (OptiPrep, CosmoBio, Japan; 60%) was diluted in MQ water to 45%, 40%, 35%, 30%, and 25%.(60) These solutions are then layered into a test tube such that the solution with highest density is at the bottom. Layers were frozen after each addition. Sample solution containing EVs was placed at the top of the tube (at RT) and ultracentrifugation (Beckman Optima L-90K Ultracentrifuge) was performed at 40,000 RPM for 16 hours at 4°C. Afterwards, 2 ml layers were separated and analyzed.

### Liquid chromatography (LC)-Mass spectroscopy (MS) and HPLC conditions

Liquid chromatography-high resolution electrospray ionization mass spectrometry (LC/MS) spectra were measured using an AB Sciex TripleTOF 4600 System (AB Sciex, Framingham, MA, USA). Chromatographic method consisted of solvent A (H_2_O with 2 mM ammonium acetate) and B (acetonitrile with 2 mM ammonium acetate), starting at 5% up to 100% of B in 10 min followed by a hold of 100% of B for 5 min, using a CAPCELL CORE C18 (2.7 μm, 3.0 × 100 mm, Osaka Soda Co. Ltd., Osaka, Japan). The method employed a flow of 0.2 ml/min, 2 μl injection volume and column temperature of 40°C. The flow rate was 0.5 ml/min and the injection volume was 2 μl. ESI-MS (R ≥ 30,000; tolerance for mass accuracy was about 5 ppm) was recorded for 15 min in the m/z region from 100 to 2000 Da.

For identifying the β-carotene in EVs, β-carotene extracted with hexane and acetone (1:1) solvent mixture from culture broth containing EVs of strain KF-425, concentrated the extract. β -carotene in the extract was measured using a Nexera Mikros System (Shimadzu Corp., Kyoto, Japan) equipped with SPD-M30A as a photodiode array detector. Chromatographic method consisted of solvent A (90% acetonitrile in H_2_O) and B (100% ethyl acetate), starting at 0% up to 50% of B in 20 min followed by a hold of 100% of B for 10 min, using a Shin-Pac Velox SP-C18 (2.7 μm, 2.1 × 150 mm, Shimadzu Corp., Kyoto, Japan). The method employed a flow of 0.2 ml/min, 2 μl injection volume and column temperature of 40°C monitoring at 450 nm.

### Spectral assignments

#### Polyphosphate

A sharp band at 1160 cm^-1^ with a broad feature at around 696 cm^-1^ indicates polyphosphate (61).

#### Polysaccharides

Two bands in the range 810 to 970 cm^-1^ are specific to glycosidic linkages in polysaccharides. C-H equatorial bending vibrations for β-type falls in the range 905–885 cm^−1^ and for α-type between 865–835 cm^-1^ (42). The broad nature of both the bands indicate complex biochemical nature of the polysaccharide cell wall of the fungus. A band at 1463 cm^-1^ corresponds to CH, CH_2_, and C-O-H deformations of the polysaccharides. Further, broad bands of overlapped peaks centered at around 1360 cm^-1^ and 1080 cm^-1^ are characteristic of polysaccharide polymer. These represents CCH/COH deformation modes and vibrations of ν(COC) glycosidic structures (62, 63).

#### Cytochrome C

A sharp band at 750 cm^-1^ can be reliably assigned to assigned to pyrrole breathing mode ν_15_ in cytochrome c (64). Sharp bands at 1584, 1312 and 1129 cm^-1^ can also be assigned to cytochrome c. The characteristic band at 1312 cm^-1^ is often regarded as the marker band for cytochrome c (in cytochrome b it appears at 1338 cm^-1^) (65).

#### Lipids

A band at 1749 cm^-1^ is due to ester the linkage in lipids. The band at 1657 cm^-1^ is an indicator of unsaturation in lipids (C=C). A relatively broad band at 1440 cm^-1^ is due to–CH_2_-scissoring mode (alkyl chain), prominent bands at 1305 cm^−1^ (CH– bending) and a broad band 1080 cm^−1^ corresponding to C–C stretching rich in gauche conformation along the carbon chain are all characteristic of lipids (36, 59).

#### Proteins

Prominent protein Raman spectral features include strong amide-I (peptide backbone vibration) at 1665 cm^−1^, C–H deformation mode at 1450 cm^−1^, tryptophan Cα–H deformation at 1340 cm^−1^, phenylalanine at 1003 cm^−1^. All these prominent features can be seen in the MCR spectral component assigned to proteins (36, 38).

#### Penicillin

Separation of Raman spectrum of penicillin presents a challenge in Raman spectroscopy method because its peaks are highly overlapped with proteins. However, using MCR, owing to its sensitivity to simultaneous variation at multiple wavenumber positions, we could separate proteins from penicillin accurately. Raman spectrum of penicillin has strong vibrational feature at 1005 cm^-1^ corresponding to benzyl group in penicillin G. A detailed study was performed earlier and thus we confirmed our assignments (38).

#### β-carotene

Three strong Raman bands are observed from β-carotene. A strong band at 1522 cm^-1^ corresponds to C=C vibrations in a conjugated alkyl chain of β-carotene. The band at 1154 cm^−1^ corresponds to C–C stretching mode coupled with C–H bending and the Raman band corresponding to C–CH_3_ rocking mode appears at 1001 cm^-1^ (39).

### Estimating composition of EVs

In order to estimate the composition of EVs, standard curves (Raman intensity vs. concentration) for benzylpenicillin potassium (in water), ß-carotene (in ethanol), and BSA (30% w/v; for proteins in EVs) were made following the standard procedures. Standard solutions of five different concentrations were prepared in each case: from 118.7 mM to 47.5 mM for benzylpenicillin potassium, 4.5 to 1.5 mM for BSA, and 24.2 μM to 8.1 μM for ß-carotene. Experimental conditions for the Raman measurements were similar to that used for Raman imaging of EVs.

## Acknowledgements

Authors thank Dr. Hiroshi Sagara and Dr. Yuji Watanabe, Institute of medical science, University of Tokyo for their help in recording TEM images. This work was supported by JSPS KAKENHI grant number JP17H06158.

## Author Contributions

A.Z.S. and H.T. conceived the idea and designed the study. A.Z.S. conducted Raman measurements, MCR-ALS data analysis, interpretation, image analysis, manuscript writing. A.Z.S, S.H. and T. N. performed penicillium culture, HPLC analysis, LC-MS analysis. A.Z.S. and N.S. performed ultracentrifugation. MCR-ALS codes written by M.A. was used in the study. A.Z.S. and H.T. has contributed to manuscript writing and discussions.

---

FIG. 1. EVs budding. a) MCR separated Raman spectral components. A full list is provided in the Fig. S2. b) Raman images of chemical constituents of P. chrysogenum filamentous body. The protrusion from the cell body is circled. An apparent location of a small opening in the cell wall is indicated with a white arrow. Size of the small opening cannot be accurately estimated (apparently <<1μm) from the image due to the limited spatial resolution of the technique. Scale bar 5 μm. c) Overlay of polysaccharide and proteins images: red color – polysaccharide and green color – proteins. d) A drawing explaining the observations. The black line indicates polysaccharide cell wall, and the protrusion is shown in green color.

FIG. 2. EVs observation and their characterization. a) white light and Raman images (overlay of β-carotene, proteins, and penicillin Raman images; for individual images see Fig. S3) of EVs from P. chrysogenum observed in the culture. Scale bar 1 μm. b) molecular constituents of the EVs imaged with Raman microspectroscopy. Molecular compositions of the EVs are characteristically different from that of filamentous body of the organism. c) estimated concentrations of molecular contents of the EVs. d) TEM images of the EVs. Scale bar 200 nm. e) Variation of penicillin content depending on the size of EVs. f) Results of HPLC analysis of concentrated supernatant containing EVs separated from 10 duplicate P. chrysogenum cultures. Elution profiles obtained by monitoring at 450 nm wavelength. UV absorption spectra of standard β-carotene solution, and β-carotene in supernatant extract (inset). h) Fluorescence image of an EVs stained with FM 4-64 membrane selective dye. Intensity profile along the line indicated in the fluorescence image is also shown. Fluorescence imaging was performed with the same Raman microscope used in the study.

FIG. 3. Biogenesis of EVs. a) Raman images of EDTA treated P. chrysogenum. Protrusions from the cell wall can be seen at several regions, and all of them have high concentration of penicillin. b) An overlay of polysaccharides and penicillin; it indicates the absence of polysaccharides in the protruding regions. c) An overlay of polysaccharide, β-carotene and penicillin: Apparently, β-carotene is present exclusively in the protrusions. Scale bar 5 μm. For the corresponding spectral components, overlay with white light image, Raman image of lipids etc. see fig. S7.

